# A model-based clustering method to detect infectious disease transmission outbreaks from sequence variation

**DOI:** 10.1101/165357

**Authors:** Rosemary M McCloskey, Art FY Poon

## Abstract

Clustering infections by genetic similarity is a popular technique for identifying potential outbreaks of infectious disease, in part because sequences are now routinely collected for clinical management of many infections. A diverse number of nonparametric clustering methods have been developed for this purpose. These methods are generally intuitive, rapid to compute, and readily scale with large data sets. However, we have found that nonparametric clustering methods can be biased towards identifying clusters of diagnosis — where individuals are sampled sooner post-infection — rather than the clusters of rapid transmission that are meant to be potential foci for public health efforts. We develop a fundamentally new approach to genetic clustering based on fitting a Markov-modulated Poisson process (MMPP), which represents the evolution of transmission rates along the tree relating different infections. We evaluated this model-based method alongside five nonparametric clustering methods using both simulated and actual HIV sequence data sets. For simulated clusters of rapid transmission, the MMPP clustering method obtained higher mean sensitivity (85%) and specificity (91%) than the nonparametric methods. When we applied these clustering methods to published HIV-1 sequences from a study cohort of men who have sex with men in Seattle, USA, we found that the MMPP method categorized about half (46%) as many individuals to clusters compared to the other methods, and that the MMPP clusters were more consistent with transmission outbreaks. This new approach to genetic clustering has significant implications for the application of pathogen sequence analysis to public health, where it is critical to robustly and accurately identify clusters for the most cost-effective deployment of resources.

## Introduction

Genetic clustering is a class of methods for reducing large sequence data sets down to groups of closely-related sequences. In the context of infectious diseases, clusters may identify infections related by a common source [1]. Additionally, genetic clusters may represent locally elevated rates of transmission, particularly when we expect a measurable number of genetic differences to accumulate within a host between transmission events. Because genetic sequencing is increaseingly a fixture of the clinical management of infections, there is growing interest in using genetic clustering as a resource for guiding public health responses in near real time [2–4]. The general motivation is that if genetic clusters define groups with higher rates of transmission, then they may facilitate a more cost-effective deployment of prevention services.

A diverse number of genetic clustering methods have been developed and applied for a broad range of bacteria and viruses including *Staphylococcus aureus* [5], *Mycobacterium tuberculosis* [6], HIV [7, 8], hepatitis C virus [9, 10] and Ebola virus [11, 12]. These clustering methods are nonparametric because the clustering criteria are based on empirical distributions, making no specific assumptions about the underlying biological processes. For instance, pairwise methods build up clusters from pairs of sequences with a genetic distance below a predefined threshold [5, 13]. Subtree methods define clusters relative to the common ancestors of sequences in the phylogenetic tree, based on quantities such as the mean branch length among the descendants of the ancestral node [14, 15]. Nonparametric methods tend to be intuitive and relatively easy to compute. We have previously observed, however, that current nonparametric clustering methods seem relatively insensitive to variation in rates of transmission [16]. Instead, these methods tend to detect variation in rates of sampling, *i.e*., the delay between infection and diagnosis.

Here we introduce a fundamentally new approach to genetic clustering for infectious diseases that is based on modelling the evolution of transmission rates along the tree. We demonstrate that our model-based (parametric) clustering method substantially outperforms a variety of nonparametric methods in recovering clusters of rapid transmission in simulated data, and we show that these differences are recapitulated in an analysis of real HIV-1 data.

## Methods

### Model

As in previous work [17, 18], we assume that the phylogenetic tree reconstructed from the genetic variation among sampled infections is similar in shape to the underlying transmission tree [19]. Hence, we assume that a branching point in the tree roughly approximates a transmission event, which is also an implicit assumption of nonparametric clustering methods where clusters are interpreted as ‘hotspots’ of rapid transmission. We model branching rates as a discrete character state that evolves along a phylogeny according to a continuous-time Markov chain. Further, we assume the branching times of the phylogeny are generated according to a Poisson process whose rate is controlled by the evolving character. Under this model, the branch lengths of the phylogeny are not independent of the character state, as typically assumed when modeling the evolution of nucleotides or amino acids.

A Cox process, or doubly-stochastic Poisson process, is an inhomogeneous Poisson process whose arrival rate *λ* (*t*) is itself a stochastic process. The Markov-modulated Poisson process (MMPP) is a special case of the Cox process where *λ* (*t*) varies according to a continuous-time Markov chain with a finite number of states. When the Markov chain is in state 1, the Poisson process has rate *λ*_1_, and so on. Following [20], we will let the Markov chain have *m* states, denote its infinitesimal generator matrix by:

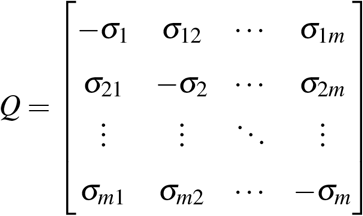

where σ_*i*_=Σ_*j≠i*_σ_*ij*_, and denote the vector of rates of the Poisson process by *λ* = [*λ*1, …, *λm*]^*T*^.

The probability density of the process producing its first arrival at time *y* in state *j*, given that it started in state *i* at time 0, is the *i j*th element of the matrix *f* (*y*) = exp((*Q −* Λ)*y*)Λ where Λ = diag(*λ*) [21]. Following [22] and [23], it is straightforward to calculate the likelihood of an observed tree under this process. Let *v* be an internal node of the *τ* other than the root, *u* be its parent, *w* and *z* be its children, and *t_v_* be the length of the branch joining *v* to its parent *u*. As in [23], define *L_i_*(*v*) to be the likelihood of the subtree rooted at *v* conditioned on the parent *u* being in state *i*. *L_i_*(*v*) is recursively defined by

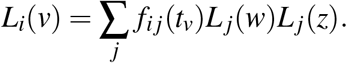

At the root of the tree, with children *w* and *z*,

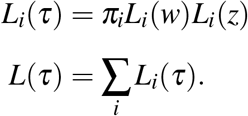

At a tip node *v* with branch length *t_v_*, it is intuitive to define the likelihood using the matrix exp((*Q −* Λ)*t_v_*), which gives the probability density of each state transition and no events occurring up to time *t_v_*. However, we found that the parameters which optimized the likelihood with this definition nearly always included one arbitrarily small rate assigned to all tips. For this reason, we simply assign *L_i_*(*v*) = 1 for all tips *v*. This approach is likely to overestimate cluster sizes due to inclusion of non-cluster individuals sampled following transmission from a cluster member, as well as individuals who are not currently part of a cluster but were in the past. In the context of viral phylogenetics, this is somewhat justified by the evidence that viruses such as HIV [24] and HCV [25] may evolve at different rates within and among hosts, implying that the branching process acting on the tips of the phylogeny may be different from the process acting on internal nodes.

To optimize the likelihood of this model, we used the covariance matrix adaptation evolution strategy [26], a black-box, derivative-free optimization algorithm. Source code implementing this model is available at https://github.com/rmcclosk/netabc.

### Data simulation

Trees were simulated using MASTER (version 5.0.2 [27]) under a susceptible-infected-removed (SIR) model of an epidemic, as described in previous work [16]. In brief, transmissions occur between infected (*I*) and susceptible individuals (*S*) at a rate *β SI*. Individuals are removed from *I* due to mortality at a rate *µ* or by becoming sampled at a rate *ψ*. The population is structured into two subpopulations with constant migration between like compartments (*S*_0_ *↔ S*_1_, *I*_0_ *↔ I*_1_) at a rate *m*. The two subpopulations comprised *S*_0_ + *I*_0_ = 9000 and *S*_1_ + *I*_1_ = 1000 individuals, respectively. Each epidemic was seeded by a single infected individual in the majority subpopulation (*I*_0_ = 1 at time 0). We simulated 100 replicate trees under three different scenarios where rates of transmission (*β*_1_) and sampling (*ψ*_1_) were varied in the minority subpopulation as follows: (1) a faster sampling rate (*ψ*_1_ *> ψ*_0_); (2) a faster transmission rate (*β*_1_ *> β*_0_); or (3) both faster sampling and transmission rates relative to the majority subpopulation. The underlying assumption is that samples from the minority subpopulation should be assigned to clusters. Infected individuals were removed due to mortality at a constant rate *µ*. The end condition for each simulation was for the tree to reach 1000 terminal branches (tips), which were subsequently filtered for tips corresponding to sampled individuals. As a result, the final number of tips was stochastic and slightly less than 1000. The simulation outputs were serialized to files in the Newick tree specification format and parsed using regular expressions in a Python script to transfer node attributes from comment strings to node labels.

In our preliminary study [16], the parameters of the model were manually adjusted until it yielded tree simulations that visually resembled the typical ‘star-like’ shape of HIV-1 molecular phylogenies (long terminal branches). In addition to expanding the number of replicate simulations, we ran a second set of simulations under different parameters to evaluate the sensitivity of our results to these settings. Specifically, we reduced the baseline (majority) transmission rate from *β*_0_ = 5 *×* 10^*−*3^ to 7.5 *×* 10^*−*4^ and reduced the sampling fraction *ψ/*(*ψ* + *µ*) — the probability that an infected individual was sampled before death — from 0.98 to 0.5 to reflect the parameter settings used in [28]. We note that this sampling fraction is very different from the sampled proportion of the infected population, since the former does not include unsampled and surviving infected individuals. Unlike [28], we increased the baseline sampling rate from *ψ*_0_ = 0.15 to 1.0 so that the latter sampled proportion increased from less than 10% to about 40%. When *ψ*_0_ = 0.15, the prevalence tended to exceed 90% at the simulation end-point where the target number of tips (*n* = 2000) was obtained. The parameter settings used in the two sets of simulations are summarized in Supporting Information (SI) Table S1. For each scenario, we discarded a small number of replicates where the epidemic failed to spread and ran additional simulations until 100 replicates were obtained.

Sequence evolution was simulated on each tree using INDELible (version 1.03) [29] with modifications to allow the user to specify the ancestral nucleotide sequence at the root. We initialized our simulations with the HXB2 *pol* reference sequence (Genbank accession K03455.1) at the root of each simulated tree. As previously described [16], the simulation parameters were calibrated to an actual alignment of HIV-1 *pol* sequences. Specifically, we used the M3 codon model with a transition rate bias *κ* = 8.0 and rate variation according to a gamma distribution (shape *α* = 1.5, rate *β* = 3) that we partitioned into 50 rate categories; a Lavalette distribution (LAV) indel model with *a* = 1.5, *M* = 4 and rate 0.001; and a scaling factor of 15.

Phylogenetic trees were reconstructed from the multiple sequence alignments using approximate maximum likelihood (FastTree2, version 2.1.10) [30] or neighbor-joining (RapidNJ, version 2.3.2) [31]. The alignments and reconstructed trees were used as inputs for five published non-parametric clustering methods — HIV-TRACE (TN93) [32], PhyloPart [33], Cluster Picker [8], subtree clustering [15], patristic distance on bootstrapped alignments [34] — and our MMPP method. Because our method requires a rooted bifurcating tree, we randomly resolved polytomies using the *multi2di* function in the R package *ape* [35] and used midpoint rooting with R package *phangorn* [36].

To measure the performance of each method, we calculated the false and true positive rates for a range of threshold parameter settings for each method, where a ‘positive’ prediction is the assignment of a sampled sequence to the minority subpopulation. For TN93 [37], we varied the genetic distance cutoff at all observed values. For Cluster Picker [8], we fixed the bootstrap threshold to 0.9 and varied the distance cutoff from 0.005 to 0.1. Similarly, we varied the maximum distance from 0.005 to 0.0125 in PhyloPart [33]. We implemented a subtree clustering method with Biopython [38], where we screened internal nodes for a bootstrap threshold of 0.9 and varied the mean branch length cutoff from 0.002 to 0.2. Lastly for the patristic method, we varied the cutoff from to 0.5 for a minimum of 80% of bootstrap replicates [34].

### Empirical data

To compare these clustering methods on an actual data set, we obtained 3102 published partial HIV-1 *pol* sequences that were previously collected in a cohort study of men who have sex with men in Seattle, USA, and analyzed for clusters of transmission [39]. We reduced the data down to a single sequence per patient (*n* = 1953) by excluding sequence records that were annotated as an additional isolate (suffixed with an underscore and integer). Next, we removed codons associated with drug resistance mutations according to the surveillance list published by Shafer and colleagues [40], using pairwise alignment of each sequence against the HXB2 *pol* reference to locate the respective codons. This step also trimmed sequence intervals that did not align to the reference. Aligned sequences that were shorter than 100 nucleotides (*n* = 22) were filtered out at this stage. We used SCUEAL [41] to predict HIV subtypes from the sequence data (maximum three recombination breakpoints) and screened for sequences categorized as subtype B excluding intraand inter-subtype recombinants (*n* = 1653). A multiple sequence alignment was generated from these data using MAFFT (version 7.305b) [42]. We reconstructed a neighbor-joining tree from this alignment with RapidNJ [31] and a maximum likelihood tree with FastTree2 [30]. Finally, we applied the different clustering methods using these trees and, if necessary, the sequence alignment, as inputs.

## Results

### Sensitivity and specificity

Epidemic dynamics were simulated from a structured birth-death model using parameter settings from [16]. We assumed that one subpopulation was larger than the other with 9000 and 1000 individuals, respectively, and that the epidemic started with a single infected individual in the larger subpopulation. This model was designed to produce genetic clusters when the epidemic moved from the larger subpopulation into the smaller (minority) subpopulation in which rates of transmission and/or sampling were elevated. Our main results are summarized in Figure 1 with respect to their true and false positive rates on 100 replicate simulations per scenario. In the first scenario, the minority subpopulation had a faster transmission rate but the same sampling rate as the majority subpopulation (‘faster transmission’). The MMPP method obtained high true positive rates (TPRs) and low false positive rates (FPRs) with means of 84.6% and 9.2%, respectively. We noticed that a small number of replicates resulted in substantially higher FPRs than the others. In these cases, we determined that the MMPP method had incorrectly assigned the ‘faster transmission’ state to the root of the tree. We subsequently determined that increasing the number of rate classes in the model enabled MMPP to correctly identify clusters with the highest rate class (data not shown). None of the nonparametric clustering methods obtained comparable TPRs or FPRs under this simulation scenario. For instance, the TN93, patristic and subtree clustering methods obtained a mean TPR of about 60% for an FPR of 30%. In contrast, Cluster Picker and PhyloPart performed poorly under this scenario, yielding cluster assignments with TPR and FPR rates that were comparable to random guessing (Figure 1).

**Figure 1:**
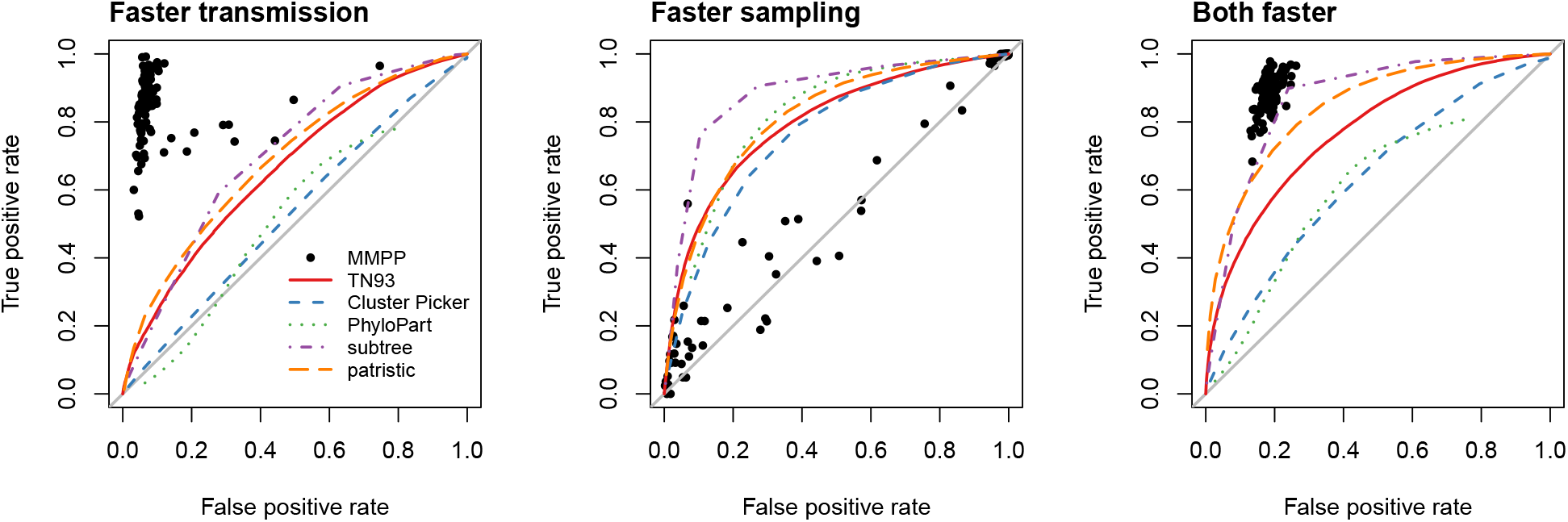
Performance of MMPP and five nonparametric clustering methods on simulated data. Sequence data were simulated under three scenarios where the minority subpopulation had (1) a faster transmission rate (left); (2) a faster sampling rate (centre), or; (3) both faster rates of transmission and sampling (right). The *x*- and *y*-axes correspond to the false and true positive rates of classifying individuals into the minority subpopulation, respectively. Each point represents the outcome when the MMPP model was applied to one of 100 replicate simulations. Each line represents the receiver-operator characteristic curve for one of the five nonparametric clustering methods (see figure legend), where different false and true positive rates were obtained by varying a threshold parameter of the method.

To illustrate the discordant results among these methods, we summarized the cluster assignments for Cluster Picker, subtree clustering and MMPP in Figure 2. In this specific example, sequences sampled from the minority subpopulation are concentrated in two groups. Clusters identified by Cluster Picker were uniformly distributed throughout the tree. In contrast, cluster assignments by the subtree clustering method were distributed less evenly with a subtree cluster coinciding accurately with one of the actual clusters. However, we found that this method was highly sensitive to the choice of mean branch length cutoff. For instance, from 0.0065 to 0.008 caused every terminal branch to be assigned to a cluster.

**Figure 2.**
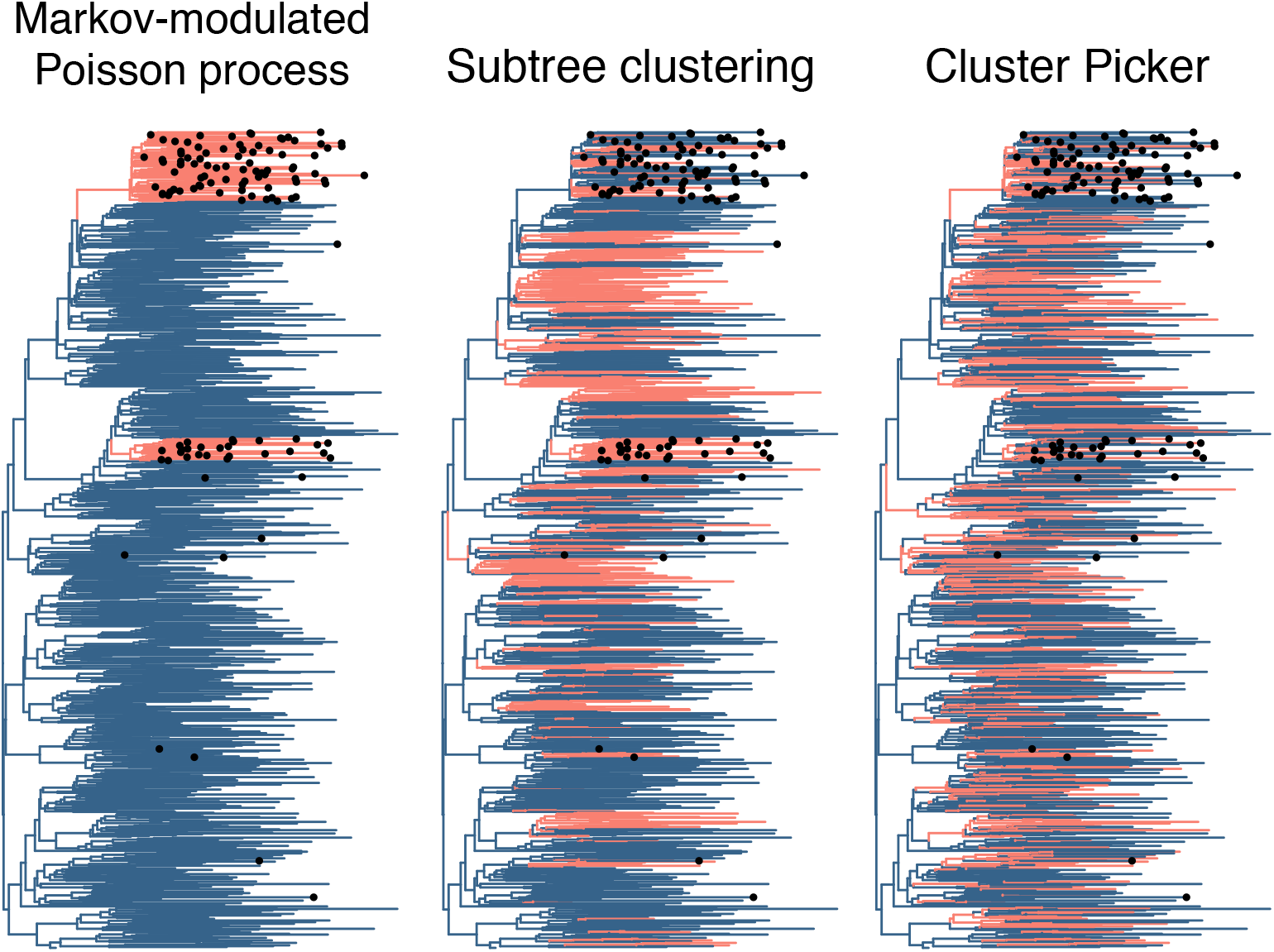
Comparison of predicted clusters and actual clusters of faster transmission. We mapped the clustering predictions from three different methods (Cluster Picker, subtree clustering and MMPP) onto one of the neighbor-joining trees reconstructed from sequence data simulated under the faster-transmission scenario. Branches are coloured light red if the method assigns that branch to a cluster, and dark blue otherwise. To colour internal branches from the nonparametric cluster assignments of tips, we used ancestral character estimation by maximum likelihood as implemented in the *ace* function in the R package *ape* [35]. The correct assignments are indicated by labeling tips with filled circles if they belong in a cluster. Cluster Picker (version 1.2.4) was run with default initial and main support thresholds (0.9) and a genetic distance threshold of 0.025. Subtree clusters were extracted from the tree with a bootstrap threshold of 90% and mean branch length threshold of 0.0065.

In the second simulation scenario, the rate of sampling was elevated in the minority subpopulation but the transmission rates were held constant (‘faster sampling’). In other words, members of the minority subpopulation were more likely to be diagnosed sooner after infection. Under this scenario the MMPP predictions were no better than a random guess with a mean TPR and FPR of 55.6% and 51.5%, respectively. This outcome was not surprising given that terminal branch lengths were excluded from maximum likelihood parameter estimation of the MMPP model. The nonparametric methods were far superior, with the best results obtained by the subtree clustering method. At a bootstrap cutoff of 90% and mean branch length cutoff of 0.006, for example, the average TPR and FPR was 78.8% and 11.9%, respectively.

Finally, the third simulation scenario combined both faster rates of sampling and transmission in the minority subpopulation (‘both faster’). The MMPP predictions attained a mean TPR and FPR of 88.7% and 18.5%, respectively. This level of performance was comparable to the subtree clustering and pairwise distance methods (patristic distance and TN93; Figure 1) at specific cutoff values. On the other hand, the Cluster Picker and PhyloPart methods both suffered worse performance under this scenario.

We obtained qualitatively similar results (SI Figure S1) when the trees were simulated under a different parameterization of the model based on [28]. Relative to the first parameter settings, the transmission rates in both subpopulations were reduced by a factor of 0.15, and the mortality rate of infected individuals was increased to equal the sampling rate in the majority subpopulation. In general, we observed higher FPR associated with the MMPP method for this set of simulations. The mean TPR and FPR of MMPP under the faster transmission scenario were 90.4% and 31.1%, respectively. To our surprise, MMPP was able to correctly identify clusters under the faster sampling scenario (82.3% TPR and 28.8% FPR), unlike the previous set of simulations. This level of performance was comparable to the subtree clustering method. We attribute this difference to the effect of lineage removal due to the elevated mortality of infected individuals. The elevated sampling rate in the minority subpopulation resulted in more complete sampling of the corresponding subtree, leading to shorter lengths of internal branches. In contrast, infected individuals were seldom removed by mortality before sampling under the previous parameterization of the model.

### Computing time

Based on our simulation analyses, the MMPP method seems to confer greater sensitivity and specificity to detect variation in transmission rates than the five nonparametric methods that we evaluated. These methods are often applied to large sequence databases with thousands or tens of thousands of records [34, 43, 44]. Furthermore, there is a growing demand for genetic clusters to be identified rapidly so that this information can be used to inform public health decisions in near real-time [4, 32]. Hence, we evaluated the average computing time require to extract clusters from simulated data sets containing approximately 1000 sequences each; the actual mean number of tips per tree was 983.3 (range 969 *−* 994). Our results, based on the times required to process five replicate data sets, are summarized in Table 1. The most time-consuming method was the patristic distance method because our default approach was to generate distances for 100 nonparametric bootstrap samples of the data [34]. Accordingly, eliminating the bootstrap sampling reduced the computing time by roughly 100-fold at the cost of sensitivity and specificity. The fastest method was TN93, which does not require reconstruction of a phylogenetic tree from sequence variation. Our MMPP method required substantially more time to compute. However, the half-minute used to process a 1000-tip tree is still an acceptable amount for near real-time monitoring.

**Table 1.**
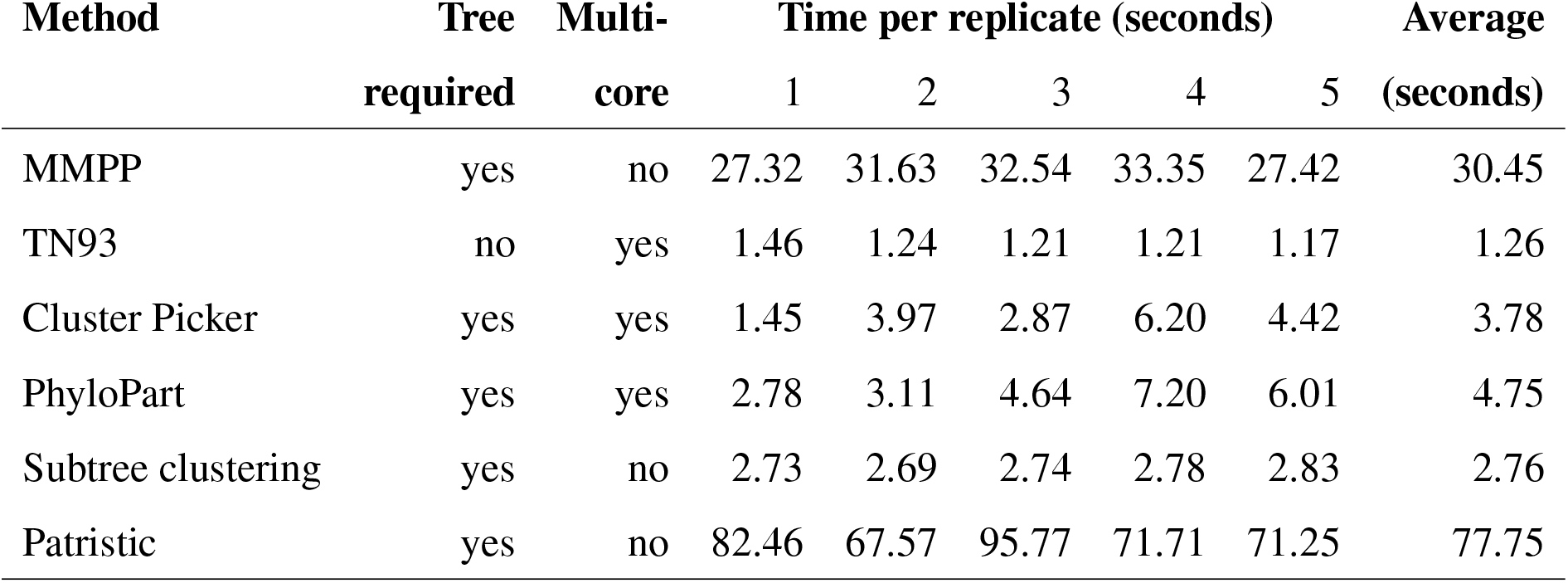
Computing time required by six clustering methods to process five different trees each relating approximately 1000 simulated sequences. All methods were evaluated on an Intel Xeon E5-1650v4 (six core) processor. If the UNIX time output implied multi-core processing and the use of multiple cores was documented for the program, then this was indicated under the heading ‘Multi-core’. None of the reported times include the time required to reconstruct phylogenetic trees from the simulated sequence data, since the specific reconstruction method used (*e.g*., maximum likelihood, neighbor joining) may vary among users. Because we observed substantial variance among repeated runs of MMPP on the same data, we reported the average of 3 runs. Times reported for TN93 include filtering the genetic distance calculations for the shortest pairwise distance per sequence.

To assess how MMPP computing time scales with the size of the tree, we generated random subsamples of 100, 200, 500 and 8000 sequences from the simulated data sets, reconstructed neighbor-joining trees for each sample, and re-ran MMPP on the resulting trees. Our results indicated that the computing time scaled non-linearly with the size of the tree: the average times were 0.82, 1.06, 2.44 and 20.68 seconds, respectively. Note that the average time to process trees averaging 983 tips each was 30.45 seconds (Table 1). These results are summarized in SI Figure S2.

### Application to real data

We obtained a published data set of *n* = 3102 HIV-1 *pol* sequences that were collected from a cohort study of men who have sex with men in Seattle, U.S. [39]. These data were reduced to one sequence per individual and then filtered for non-recombinant subtype B sequences (*n* = 1653, see Methods). We reconstructed a maximum likelihood phylogeny from these sequences as the primary input for the different clustering methods. First, we discovered that the MMPP method tended to assign faster branching rates throughout the base of the tree, including the root node. A lineages-through-time plot of the tree (SI Figure S3) was consistent with a period of population-level exponential growth in the first half of the tree, which may confound the MMPP model from detecting other sources of branching rate variation over time. Based on our experience with simulated cases where the faster of two rate classes was sometimes incorrectly assigned to the root of the tree, we increased the number of rate classes to 3 and grouped lineages assigned to the fastest rate class as putative clusters. The resulting clusters are summarized in Figure 3, alongside the clusters predicted by two nonparametric methods (TN93 and subtree clustering). Our MMPP program required 43.6 and 28.7 seconds to analyze the ML tree assuming two and three rate classes, respectively. Without adequate information to exactly reproduce the clusters reported by the source study [39], we tuned each nonparametric clustering method until the number of individuals assigned to clusters was similar to the number reported in that study (*n* = 168 individuals in 72 clusters).

**Figure 3:**
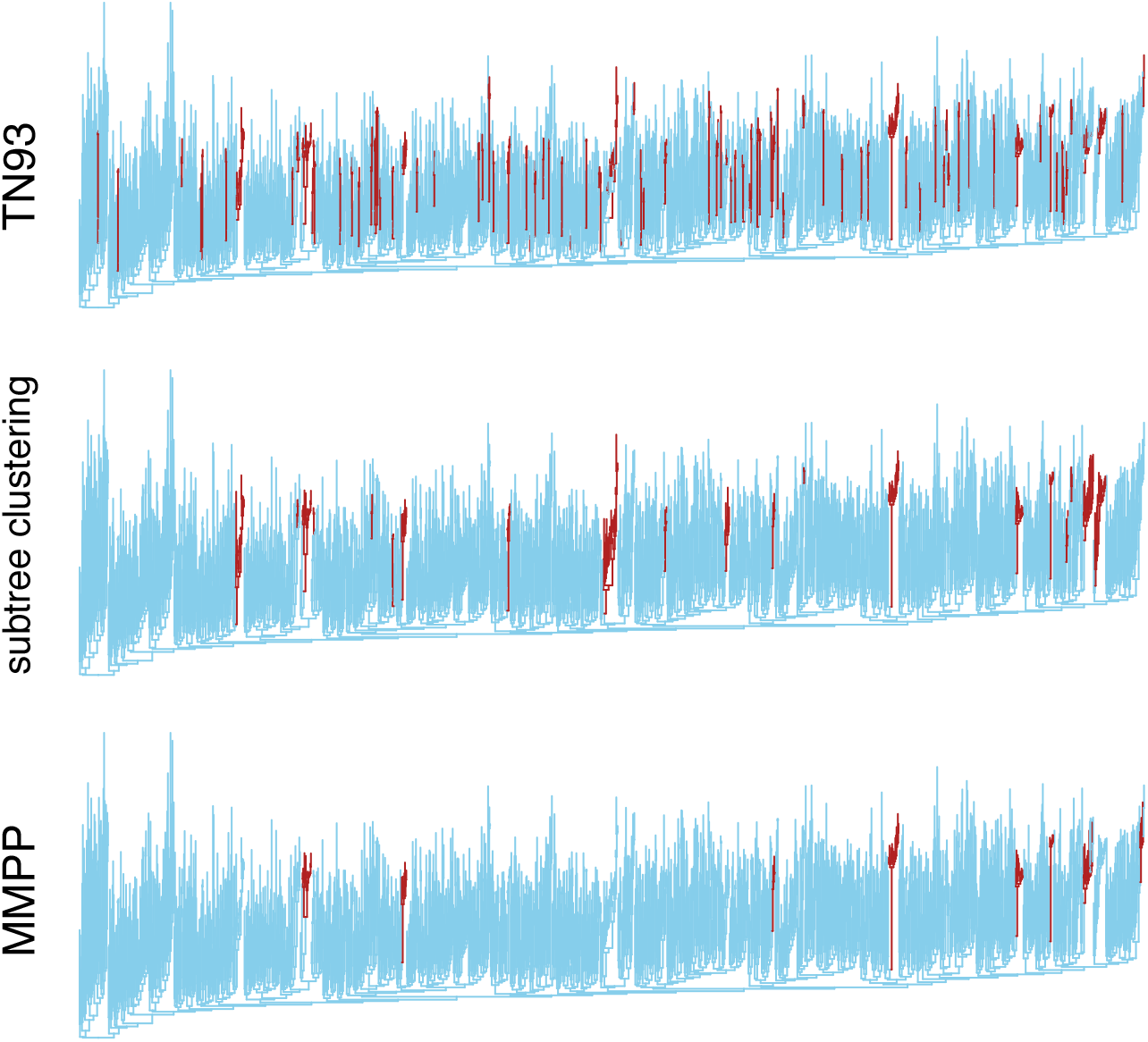
Comparison of three different clustering methods applied to a phylogeny reconstructed from real HIV data. The phylogeny was reconstructed by maximum likelihood from a published data set of HIV-1 subtype B *pol* sequences sampled from a study cohort of men who have sex with men in Seattle, U.S. [39]. We adjusted the TN93 and subtree clustering methods until the number of individuals assigned to clusters was similar to the number reported in the original study. Branches assigned to clusters are highlighted in dark red.

The most apparent difference between the nonparametric methods and the MMPP results, which are summarized in Figure 3, was the number of individuals assigned to clusters. Our MMPP model assigned only 78 individuals (150 branches in the tree) to 8 clusters, substantially fewer than the number reported in [39]. We note that the number of individuals in nonparametric clusters is determined by the subjective selection of clustering criteria. To match the number reported by [39], the TN93 method required a distance cutoff of 0.006, and our implementation of the subtree method used a mean branch length cutoff of 0.0065 and a bootstrap cutoff of 0.9. Based on the assignment of branches to clusters, MMPP was more concordant with the subtree method (Cohen’s *κ* = 0.573) than TN93 (*κ* = 0.341), where *κ* = 1 indicates complete agreement. Within clusters identified by the subtree method but not by MMPP, terminal branch lengths (*n* = 92) were significantly shorter (Wilcoxon test, *P <* 10^-12^) and internal branch lengths (*n* = 64) were no different (*P* = 0.24) from their respective distributions in the entire tree. Similarly, terminal branches in TN93 clusters not identified by MMPP (*n* = 124) were significantly shorter (*P <* 10^-12^) but the internal branches (*n* = 31) were only marginally shorter (*P* = 0.051). In contrast, internal branches were significantly shorter in MMPP clusters (*P* = 5.98 *×* 10^-7^). These results suggest that many of the nonparametric clusters may be caused by variation in sampling rates; however, we cannot ascertain the extent of this effect without additional information such as estimated dates of infection.

## Discussion

Here we have formulated and tested a model-based (parametric) method for clustering genetic sequences that have been sampled from an infectious disease epidemic. A potential advantage of a model-based approach is that we can focus on the parameter of interest — namely, the variation in transmission rates over time that may be an indicator of outbreaks. Our approach is conceptually similar to the lineages-over-time model proposed by Holmes and colleagues [17]; however, we assume that branching rates evolve along branches of the tree, such that discordant rates may occur on contemporaneous branches on different parts of the tree. The MMPP model is also closely related to models of biological speciation where rates of speciation are determined by a single evolving character state, *e.g*., [45, 46]. Specifically, our model can be interpreted as a special case of the multitype speciation-extinction model [47] where extinction events are censored from the model. Our simulation results indicate that the interpretation of MMPP clusters with respect to sampling rates is dependent on the probability that a genetic sequence of the pathogen is sampled from an infected individual before their death. If most infected individuals are eventually sampled, then we predict that MMPP clusters will largely be determined by variation in transmission rates. If a substantial fraction of infected individuals are never sampled before their death, however, then the MMPP clusters may be caused by variation in either rates of transmission or sampling. Nevertheless, we submit that this confounding is preferable to being unable to detect clusters of transmission at all (Figure S1).

When interpreting clusters produced under the MMPP model, we assume that the variation in the rates of branching events in the phylogeny is a sufficient approximation of variation in transmission rates over time. The same assumption is implicit when nonparametric genetic clusters are interpreted as putative transmission outbreaks. There are several reasons why transmission events are likely to map to locations of the virus phylogeny other than the branching points, such as incomplete lineage sorting within hosts [48] and incomplete sampling of the infected population [49]. Another issue that is unique to model-based clustering is the problem of model misspecification. The present MMPP model assumes that rates of transition between branching rates are constant over time. For instance, fitting the MMPP model with two rate classes to the actual HIV-1 data set resulted in a majority of branches assigned to the faster rate class, including the root of the tree. This was likely caused by an early period of exponential growth in the epidemic (Figure S3), which induces lineage-independent variation in branching rates over time. Furthermore, the spread of an epidemic through a socially and spatially-structured host population is problematic for the MMPP model, which assumes that the evolution of branching rates is a memoryless process. These issues identify directions for further work in this new class of genetic clustering methods for infectious disease outbreaks.

Genetic clustering can be an important resource for retrospective epidemiological investigations [50] and may eventually play a central role in the near-real time monitoring and prediction of infectious disease outbreaks [3, 4]. However, the growing popularity of applying genetic clustering to detect outbreaks of transmission needs to be tempered with greater skepticism about the underlying methods [51]. Our results suggest that many clustering methods are potentially misdirecting public health efforts away from groups suffering from higher rates of transmission, and towards groups where new infections were diagnosed sooner than the population average. As we have shown with our analysis of the MMPP method, bringing new approaches to the clustering problem may provide a more complete picture, in combination with current methods, about the recent history of an epidemic.

## Acknowledgements

We wish to acknowledge Simon D. W. Frost for his insightful and helpful feedback on an earlier version of this manuscript.

